# Large-FOV 3D localization microscopy by spatially variant point spread function generation

**DOI:** 10.1101/2023.07.30.551150

**Authors:** Dafei Xiao, Reut Kedem Orange, Nadav Opatovski, Amit Parizat, Elias Nehme, Onit Alalouf, Yoav Shechtman

## Abstract

Accurate characterization of the microscopic point spread function (PSF) is crucial for achieving high-performance localization microscopy (LM). Traditionally, LM assumes a spatially-invariant PSF to simplify the modeling of the imaging system. However, for large fields of view (FOV) imaging, it becomes important to account for the spatially variant nature of the PSF. In this work, we propose an accurate and fast principal component analysis (PCA)-based field-dependent 3D PSF generator (PPG3D) and localizer for LM. Through simulations and experimental 3D single molecule localization microscopy (SMLM), we demonstrate the effectiveness of PPG3D, enabling super-resolution imaging of mitochondria and microtubules with high fidelity over a large FOV. A comparison of PPG3D with three other shift-invariant and shift-variant PSF generators for 3D LM reveals a three-fold improvement in accuracy and an operation speed approximately one hundred times faster. Given its user-friendliness and conciseness, we believe that PPG3D holds great potential for widespread application in SMLM and other imaging modalities.

## 1. Introduction

Localization microscopy (LM) is a powerful imaging modality both for super-resolution imaging^1,2^ and for particle tracking^3^. Imaging in these modalities is based on determining the positions of point sources or emitters at a sub-diffraction precision.

Emitters from a flat sample can be localized to yield a 2D image, however, in the case of a volumetric sample, 3D LM is needed. One approach for 3D LM is PSF engineering, which intentionally induces aberrations at the pupil plane of an imaging system to yield an informative depth-dependent PSF, e.g., astigmatism^4^, double-helix^5,6^, or tetrapod^7^. For the decoding part, common localization algorithms include maximum likelihood estimation^8–10^ and more recently, deep-learning-based algorithms, e.g., DeepSTORM3D^11^ and DECODE^12^, which have demonstrated excellent performance when dealing with PSF overlap and high emitter-density. Considering the influence of field position on localization (field-dependent localization), recent work^13^ proposes FD-DeepLoc by combining DECODE with CoorConv^14^ to both address the field dependence and estimate emitter positions. Importantly, all estimators rely on some forward model, a PSF generator, i.e. a continuous-domain “dictionary” that maps an emitter’s 3D spatial position to a PSF. The performance of LM strongly depends on the accuracy of this PSF generator^15^.

The common assumption in LM is that the system is shift-invariant, which yields shift-invariant PSF generators. In its basic form, a PSF generator receives as input the optical system parameters and a 3D position and outputs the expected image on the camera of an emitter in that position. The generator is considered shift-invariant if it is independent of the global transverse position of the emitter, which simplifies the PSF’s spatial dependence to be depth-only.

Shift-invariant PSF characterization is typically based on a physical model^16,17^, with additional experimental input to account for unavoidable aberrations and deviations from the ideal models. A commonly used analytical model for the 3D microscopic PSF is based on the principle that the PSF can be obtained by taking the absolute value squared of the Fourier transform of the field at the Back Focal Plane (BFP)^18^; emitter defocus manifests as an approximately quadratic phase added to the BFP field. The experimental input can be acquired through a calibration experiment, where the PSF is measured at various known axial positions. Then, combining the calibration with the analytical model involves solving a phase-retrieval problem^19^ to determine the BFP phase profile that aligns with the experimental calibration measurement. This BFP phase profile can be projected onto Zernike polynomials, for computational efficiency, to yield Zernike pupil phase retrieval (ZPPR)^20,21^. For improved performance, a pixel-wise pupil representation can be used, at a cost of more optimization parameters, yielding VIPR (short for vectorial implementation of phase retrieval)^18^. In addition to these model-based approaches, model-free generators utilize polynomial B-spline^22^, C-spline^23^ basis, and Zernike moments/basis^24^ to represent the depth-dependent PSF and yield PSF interpolation.

The shift-invariant assumption, however, deviates from reality in several cases. First, objectives, especially high-numerical-aperture ones widely used in SMLM, exhibit inherent field-dependent aberrations that vary with transverse positions^21,25^. Second, PSF engineering in 3D LM introduces additional optical elements that inevitably add some field dependence, e.g. due to some displacement of the phase element from the BFP. Third, field dependence becomes more prominent as FOV increases^13,26,27^, which is increasingly relevant in high-throughput imaging scenarios. Therefore, for 3D LM at high fidelity over a large FOV, the development of a PSF generator that considers both depth and field dependence would be highly beneficial.

In the context of LM, to the best of our knowledge, the only PSF generator that consider field dependence are based on ZPPR. Shift-variant ZPPR algorithms^13,21^ rely on calibration measurements at many field positions to retrieve a Zernike-polynomial-based pupil phase for each position. Then, interpolation of the Zernike basis coefficients is implemented to obtain the pupil phase corresponding to any desired field positions. In other fields, e.g., astronomy^28^ and computational volumetric imaging^29,30^, field dependence has been characterized through truncated singular value decomposition (TSVD) of calibrated PSFs and interpolation of the decomposition coefficients.

Here, we extend the PSF decomposition and interpolation concept and propose a fast and accurate spatially variant PSF generator in 3D (PPG3D) specifically designed for SMLM. PPG3D is a continuous-domain PSF model which implements principal component analysis (PCA) of calibrated PSFs, conducts local interpolation of those principal component (PC) coefficients, and then generates the PSF at any target position through backward space projection. Comparison of PPG3D with three other commonly-used PSF generators in LM demonstrates improvements of over three times in accuracy and around a hundred times in computation speed. In its application to SMLM, we combine PPG3D with our localization estimator FOV-dependent DeepSTORM3D, customized from one of the state-of-art localization estimators, DeepSTORM3D^11^, and achieve 3D super-resolution imaging of mitochondria and microtubules with high fidelity over a large FOV.

## 2. Results

### 2.1. PPG3D

#### PPG3D formation

PPG3D is a PSF interpolator built upon calibrated PSFs at known spatial positions. Direct pixel-by-pixel interpolation of the measured PSF is prone to failure due to sensitivity to noise and ill-posedness. To address this issue, we employ PCA of the measured PSFs (Tetrapod PSF engineering in Fig. 1 (a)) and perform 3D interpolation in a lower dimensional space (Fig. 1 (b)). Because lateral-position-dependence is more moderate compared to depth dependence for PSF of SMLM, we separate the 3D interpolation into two 2D lateral interpolations and one 1D axial interpolation. Furthermore, we empirically find that a PSF in LM can be adequately represented by its surrounding PSFs. As a result, we employ local interpolation, specifically three lateral positions and two axial positions (Fig. 1 (b)), to improve the operation speed.

**Figure 1.**
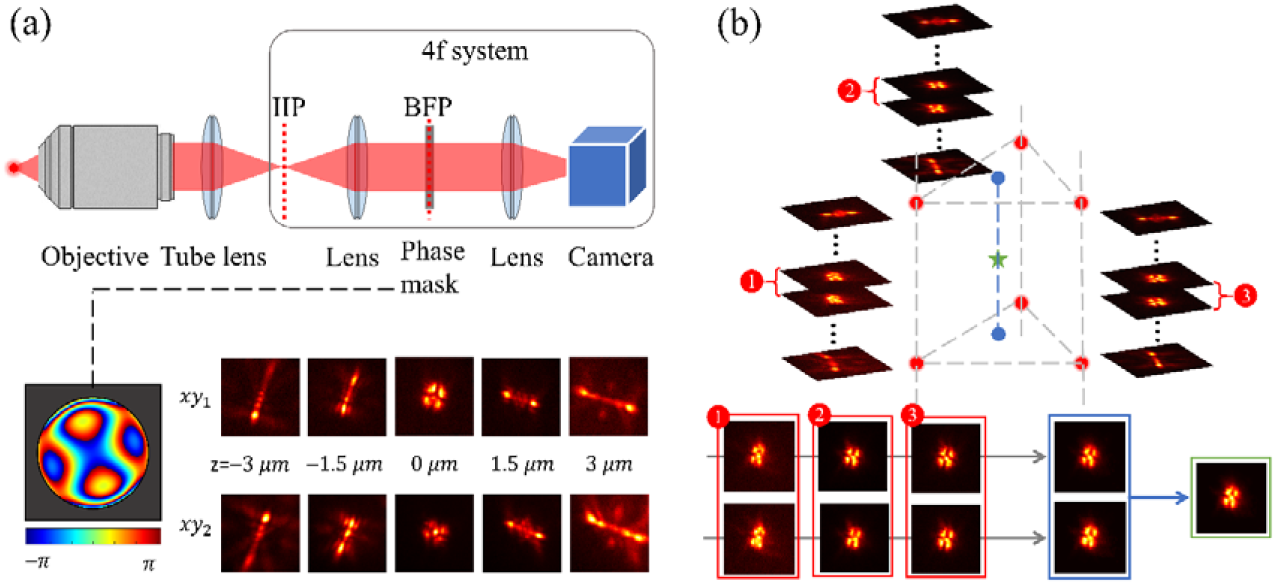
Schematic of PSF engineering and PPG3D. (a) PSF engineering with a Tetrapod phase mask at the back focal plane (BFP) and experimentally measured engineered PSFs at two field positions (white squares in Fig. 2 (a)) exemplifying field dependence. IIP denotes intermediate imaging plane. (b) PSF at a desired position (green star) is interpolated using six measured PSFs at nearby calibration positions (red dots). Each lateral position corresponds to a z-stack of depth-dependent Tetrapod PSF. The bottom flowchart shows two separate steps in PPG3D: lateral (gray arrows) and axial interpolations (blue arrow).

To generate a PSF at any spatial position, we first search for three lateral calibration positions close to the target coordinate (Fig. 1 (b)) according to a criterion of even distribution (section 3 of SI), at two axial planes above and below the target axial position, such that a total of six PSFs are chosen. Next, PCA of selected PSFs at each axial plane is conducted and lateral 2D interpolations regarding PC coefficients are performed to address field dependence, which yields two PSFs, located at the blue dots in Fig. 1 (b). The following step is PCA of those two laterally interpolated PSFs and a 1D interpolation of the PC coefficients, followed by a transformation back to image space, to yield the final target PSF. More details of the implementation are in section 3 of the SI and field dependce quantification is presented in section 5 of the SI.

#### Comparison of PPG3D with PSF generators in LM

The comparison is performed upon a calibration dataset (151 lateral positions by 29 axial positions) obtained from an imaging system with tetapod PSF engineering (see imaging system #1 of Table S1 in SI). Each cropped PSF has a cropped size of 81×81 pixels (see PSF cropping details in section 2 of SI). We divide the comparison into the shift-invariant case, i.e. axial-only interpolation, and the shift-variant case, i.e. axial and lateral interpolation. The criteria for assessing different generators include correlation coefficient (CC), root mean square error (RMSE), comparing measured PSFs to generated ones, as well as runtime.

We first compare PPG3D’s shift-invariant mode with three shift-invariant PSF generators: model-based VIPR and ZPPR, and a model-free Spline interpolator. At the field position P0 in Fig. 2(a), near the center of the FOV, PSFs are measured at a series of axial positions (Fig. 2 (b) horizontal axis) which are divided into a “seen” group and an “unseen” group (marked by grid lines in Fig. 2 (b)). The former is set as input for PSF generation and the latter is used for testing. The details of implementing VIPR and ZPPR are shown in section 4 of SI. Firstly, PPG3D exhibits larger CC and lower RMSE, compared with others, at “unseen” axial positions (Fig. 2(b)), which is also visually demonstrated by generated PSFs at a test position (black dot line in Fig. 2 (b)) in Fig. 2 (c). Secondly, PSFs generated at “seen” positions by PPG3D are nearly identical to the measured ones, while all other generators exhibit some characterization errors. Thirdly, among the two model-based generators, VIPR outperforms ZPPR, thanks to its pixel-wise optimization. Finally, in terms of runtime measured through the codes we can access (MATLAB codes for VIPR and ZPPR, Python codes for Spline and our PPG3D), (Fig. 2 (d)), PPG3D is around 100 times faster than the second fastest algorithm, which is VIPR.

**Fig 2.**
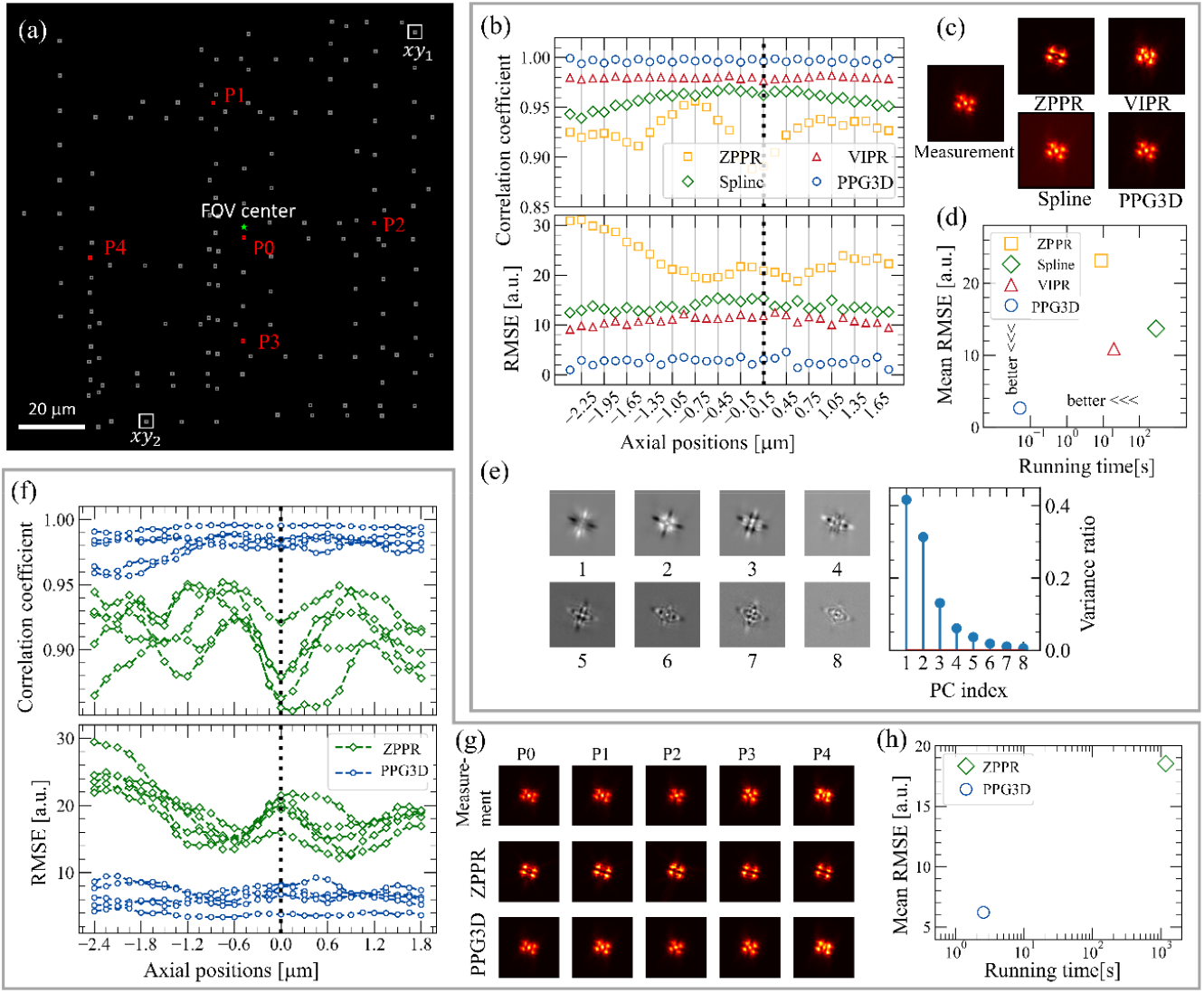
Comparison of PPG3D with three PSF generators in LM. (a) A FOV with 151 calibration field positions (gray and red dots). The comparison is divided into shift-invariant mode (b)-(e) considering point P0 in (a), and shift-variant mode (f)-(h) where the red dots are set as unseen test positions. (b) CC and RMSE assessment of ZPPR, VIPR, Spline, and PPG3D. The axial positions with grid lines are set as unseen test positions. (c) Generated PSFs at the black dot line of (b). (d) RMSE-time comprehensive assessment. (e) Principal components used in PPG3D and their variance ratios. (f) CC and RMSE assessment of shift-variant ZPPR and PPG3D at the five test positions. (g) Generated PSFs at the lateral test positions and one axial position (black dotted line in (f)). (h) RMSE-time comprehensive assessment.

Next, we compare PPG3D’s shift-variant mode with shift variant ZPPR. Among all the calibrated field positions, we choose 5 as test positions (red dots in Fig. 2 (a)). At all 5-by-29 test spatial positions, PPG3D performs better in both CC and RMSE than ZPPR (Figure 2 (f). The implementation details of shift-variant ZPPR are presented in Fig. S3 of SI). Specifically, ZPPR fails to characterize the central bright spot of measured PSFs (Figure 2 (g)), caused by unmodulated light at the BFP, which often occurs due to some mismatch between the phase-mask size and the actual light circle at the BFP. Meanwhile, PPG3D is hundreds of times faster than shift-variant ZPPR, which is important for applications such as online training of deep-learning-based localization algorithms.

### 2.2 High-accuracy large-FOV 3D SMLM

#### A field-dependent localization estimator: FOV-dependent DeepSTORM3D

We construct a complete field-dependent 3D SMLM localizer, by combining PPG3D with DeepSTORM3D^11^, a deep learning-based algorithm for 3D SMLM localization, generating a large-FOV field-dependent localization estimator: FOV-dependent-DeepSTORM3D. This is based on the addition of a CoordConv^14^ to all the feature detection layers of DeepSTORM3D (Fig. 3). To overcome the computational challenges from the large FOV reconstruction, we employ a segmentation and consolidation scheme which involves cropping the complete image into sub-images for localization and subsequently stitching together the localization results. Notably, PPG3D can be combined with localizers other than DeepSTORM3D, e.g. DECODE^12^, which is used in FD-DeepLoc^13^.

**Fig. 3.**
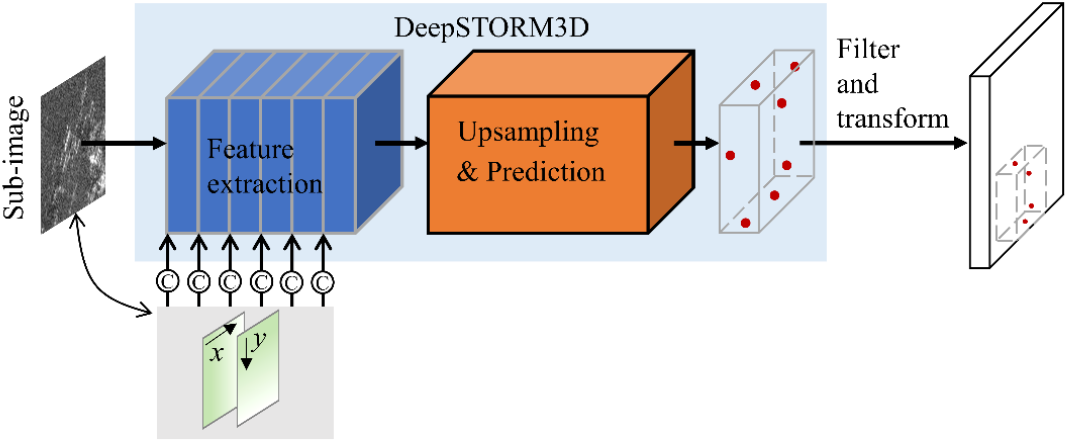
FOV-dependent DeepSTORM3D. Based on DeepSTORM3D, x and y maps corresponding to the cropped sub-image are fed to all six feature extraction layer to yield local localizations, which are then filterd and transformed into final global localizations.

In the training phase, training images consist of generated sub-images with random PSFs placed away from the sub-image edges to prevent PSF truncation. During the inference phase, we divide each experimental image into sub-images with some overlap. These sub-images are then fed individually into the network to obtain localizations, and the localizations from all sub-images are fused to generate the complete global localization map for the frame. To ensure accuracy, unreliable localizations near the edges of each sub-image are excluded, focusing only on the central valid area. Notably, this exclusion does not result in localization loss due to our precise cropping overlap design (Fig. S7 of SI).

#### Network training and inference on simulated data

Using PPG3D, we generate field dependent training data for FOV-dependent DeepSTORM3D, which divides the whole FOV into 11-by-11 patches with some overlapping (Fig. S7 in SI). The bottom-left 6-by-6 patches (Fig. S7 in SI), around a quarter of the whole calibrated FOV, are used to demonstrate our concept in simulation. For comparison, we also train a standard DeepSTORM3D with the same training images which, however, are from the shift-invariant mode of PPG3D. First, we simulate 240 random emitters and test localization accuracy according to RMSE. Note that both networks, in the prediction stage, use a thresholdregarding prediction confidence. Because the same threshold for both networks yields different number of localizations in the same image, we set a series of thresholds and show a plot of localization RMSE to the number of predictions. Furthermore, for each threshold, we repeat the inference ten times with different random emitters. The lateral and axial RMSE between the predicted localizations and ground truth are calculated for assessment (Fig. 4 (a) and (b)). On average, FOV-dependent DeepSTORM3D improves the lateral and axial localization accuracy by 36% and 30% respectively.

**Fig. 4.**
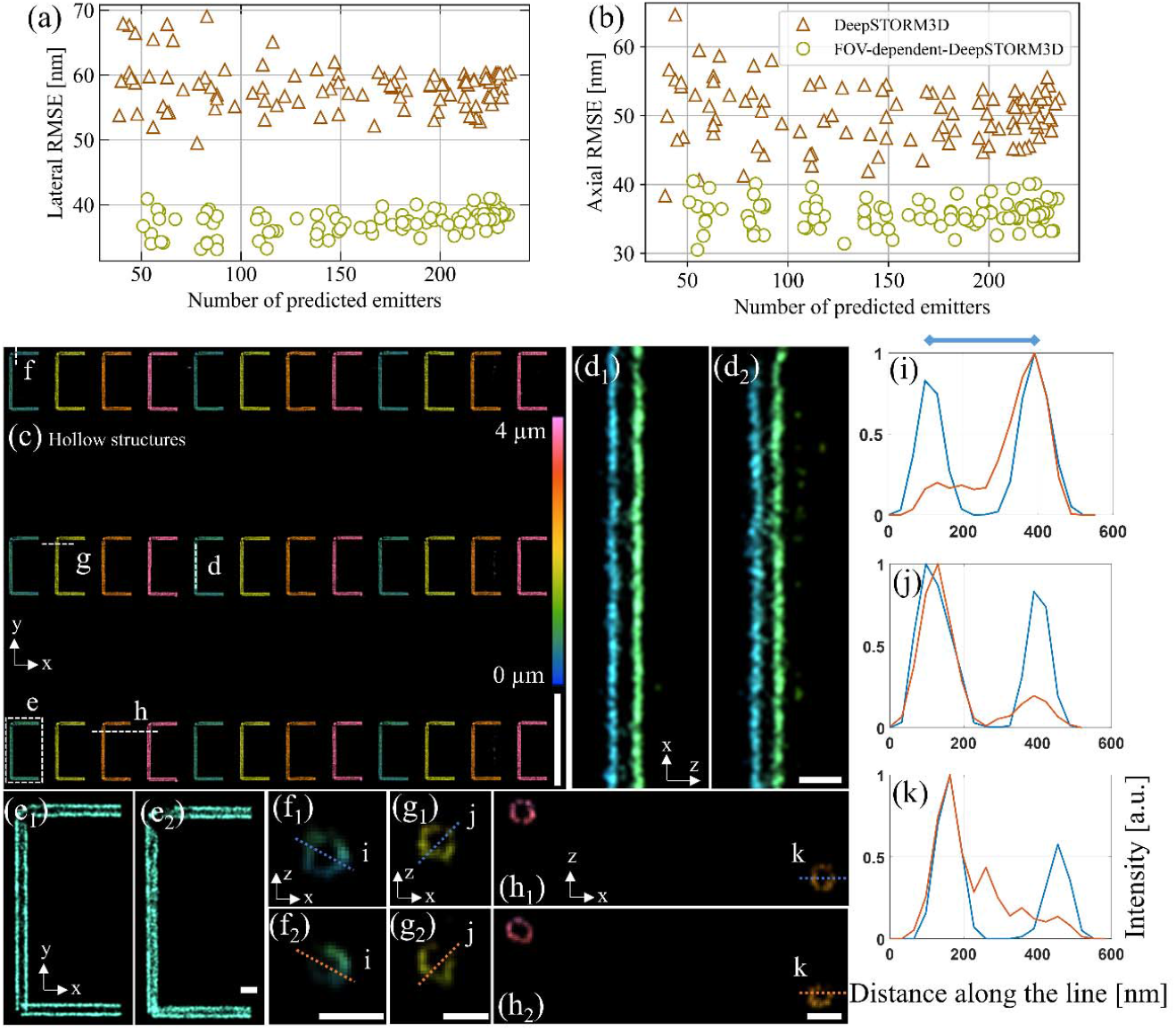
Comparison of FOV-dependent DeepSTORM3D (subscript 1) to DeepSTORM3D (subscript 2) in simulation. (a) Lateral RMSE of predictions with respect to the number of emitters detected by both networks. (b) Same comparison regarding axial RMSE. (c) Reconstructed 3D dotted tubular “C” structure by FOV-dependent DeepSTORM3D. Scale bar: 10 μm. The cross sections along the lines d, f, g, and h are in (d), (f), (g), and (h), and a zoom-in view of e at z= 6.175 μm is shown in (e). Scale bar: 0.5 μm. The red arrow represents z-axis. The intensity profile along i in (f), j in (g), and k in (h) are shown in (i), (j), (k). The blue line on the top of (i) shows the true diameter of the tube, 300 nm.

Secondly, we create a 3D structure with 9 units of “CCCC”. Each unit is composed of 16 dotted tubules, each of which has a diameter of 300 nm, length of 3 μm, and consists of 3200 emitters (details in section 7.1 of SI). Simulating the emitter blinking in a SMLM experiment, we predict the structure using 4000 frames of simulated images. When processing those images through both networks, we ensure the approximately same number of interferences through filters in ThunderSTORM^31^ (section 7.2 of SI). As the reconstuction (Fig. 4 (c-k)) shows, FOV-dependent DeepSTORM3D (subscript “1” in the Fig. 4) outperforms standard DeepSTORM3D (subscript “2”) and yields super-resolution reconstruction with higher fidelity. Specifically, Fig. 4 (d_2_) shows noisier and deformed reconstructed tubules, which is imporved and corrected in Fig. 4 (d_1_). Similarly, Fig. 4 (e_1_) presents more clear hollow tubule structure compared with Fig. 3(e_1_). More detailed quantification comparisons regarding hollow structure (Fig. 4 (d-h) and intensity profiles in Fig. 4 (i-k)) demonstrates that FOV-dependent DeepSTORM3D helps correct misestimations caused by field dependence. An additional simulation of a sinosoidal surface is shown in Fig. S9 of SI.

#### Experimental demonstration: super-resolution imaging of mitochondria and microtubules

We perform 3D STORM using the Tetrapod PSF (see imaging system #2 in Table S1 in SI). The experiment consists of imaging a fluorescently labeled samples as the fluorophores blink in time, localizing them, and reconstructing a super-resolved image using either FOV-dependent DeepSTORM3D or DeepSTORM3D. See experimental details in the Methods section. We first image mitochondria, over a complete FOV of 178-by-178 μm; in total, 52261 frames (see one example frame after background subtraction in Fig. 5 and blinking movie S1) were processed; and ∼12.7 million valid localizations were found. Comparing the localization results (Fig. 6) from FOV-dependent DeepSTORM3D (subscript “1”) and DeepSTORM3D (subscript “2”), the improvement is noticeable. Specifically, Fig. 6 (d_1_) shows a shaper pink and orange structure compared with Fig. 6 (d_2_). Also, Fig. 6 (e_1_) and (f_1_) outperform their counterparts considering the clarity and cleanness of the microstructures. Furthermore, the cross sections (g-j), as well as the intensity profiles in (k-n), show that cleaner, more solid, and more complete hole structures can be detected when considering field dependence.

**Fig. 5.**
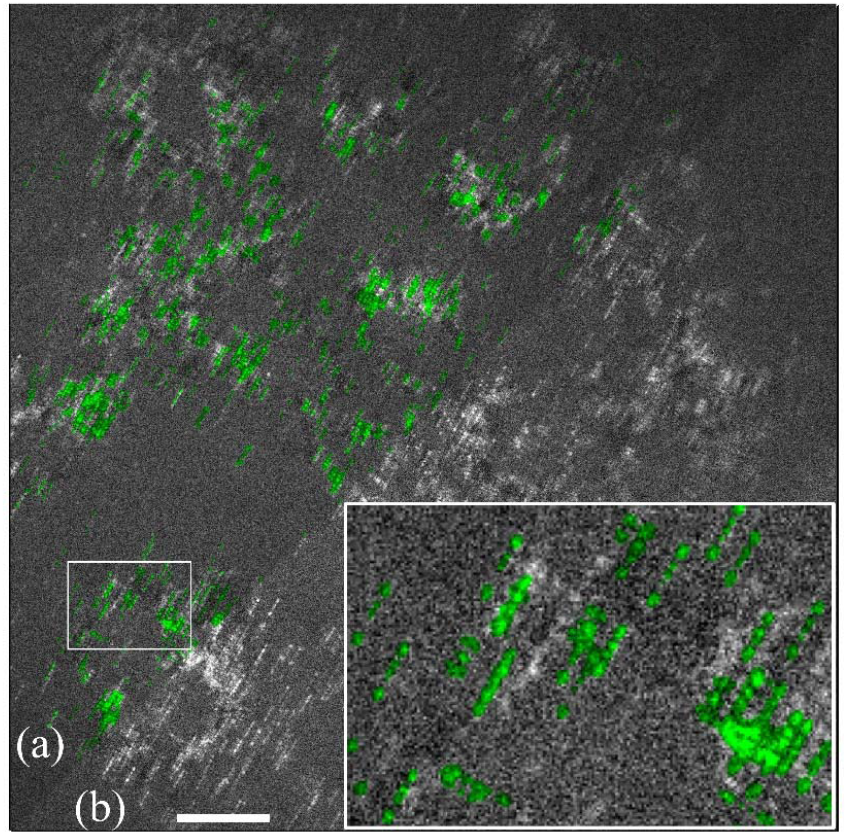
Demonstration of one representative experimental frame (b) and the rendered image (a) which is an overlay, upon this frame, consisting of the reconstructed PSFs according to network inference. Scale bar = 20 μm. The bottom-right is the zoom-in view of the white square at the diagonal.

**Fig. 6.**
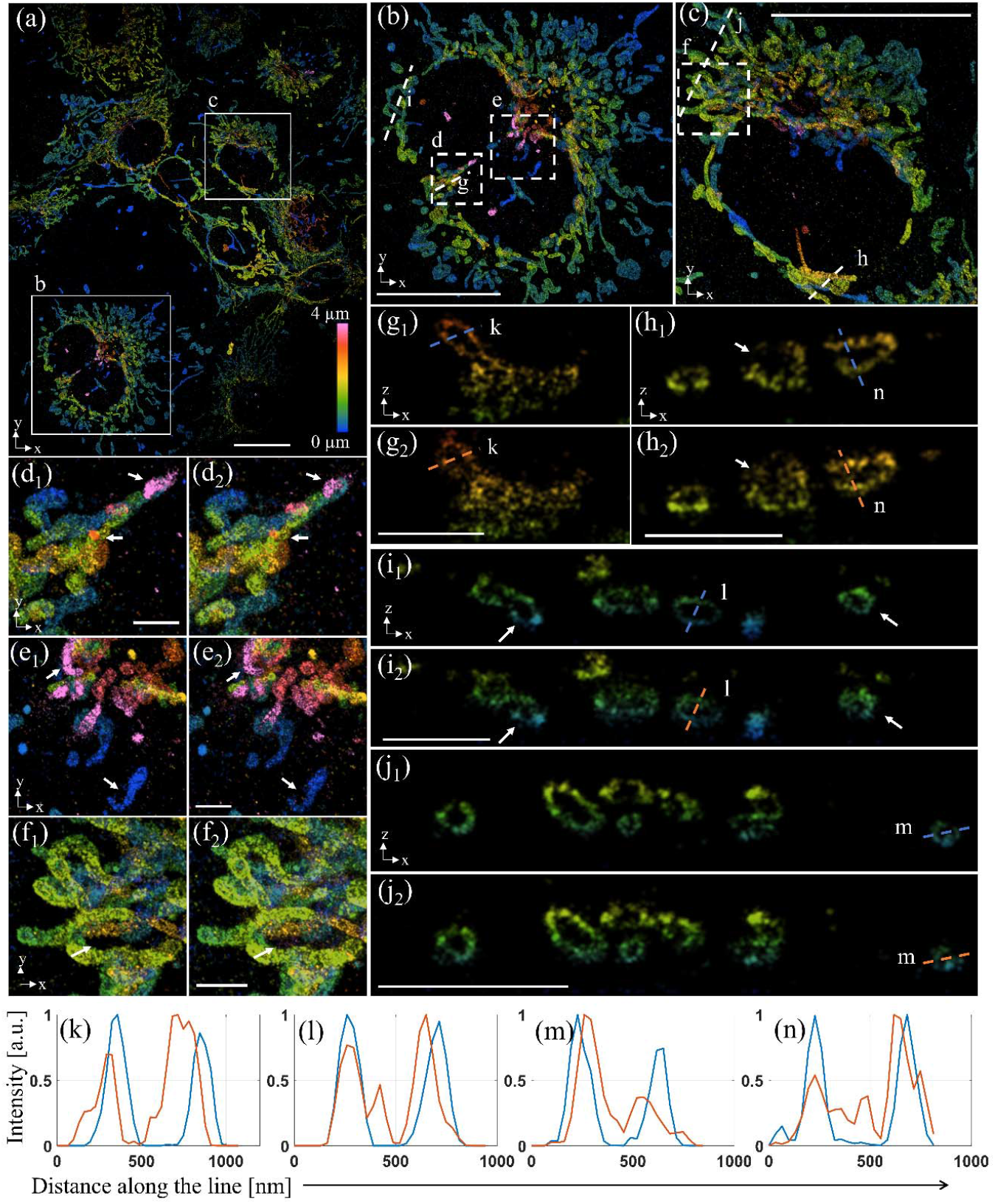
Comparison of 3D super-resolution image of mitochondria between FOV-dependent DeepSTORM3D (subscript “1”) and DeepSTORM3D (subscript “2”). (a), Super-resolved mitochondria imaging (maximum projection) in a 127-by-160 μm FOV through FOV-dependent DeepSTORM3D. Scale bar: 20 μm. The zoom-in views of square regions b and c are shown in (b), (c). Scale bar: 20 μm. The comparison of regions d and e in (b), and f in (c) are shown in (d), (e), and (f) respectively. The white arrows indicate some details worthy of comparison. Scale bar: 2 μm. The cross sections along lines g and i in (b), and j and h in (c), are shown in (g), (i), (j), and (h) with the red arrows indicating z axis. Scale bar: 2 μm. The intensity profiles along lines in those cross sections are shown in (k-n). The blue and orange lines correspond to FOV-dependent DeepSTORM3D and DeepSTORM3D, respectively.

Next, we also image a microtubule sample (Fig. 7). A total of 52322 fames of experimental images with dense emitters were processed and ∼11.5 million valid localizations were found for both networks. Because field-dependent aberrations become more prominent with more defocus (closer to 0 nm and 4000 nm in our case), we focus on axial positions of ∼100 nm and 4000 nm and show the corresponding reconstructed tubule structures from FOV-dependent-DeepSTORM3D (Fig. 7 (b_1_) and (c_1_)) and DeepSTORM3D (Fig. 7 (b_2_) and (c_2_)). As the comparisons show, field dependence consideration enables a more complete and solid reconstruction. An additional experimental demonstration with less dense cells and finer microtubule details is shown in Fig. S10. of SI.

**Fig. 7.**
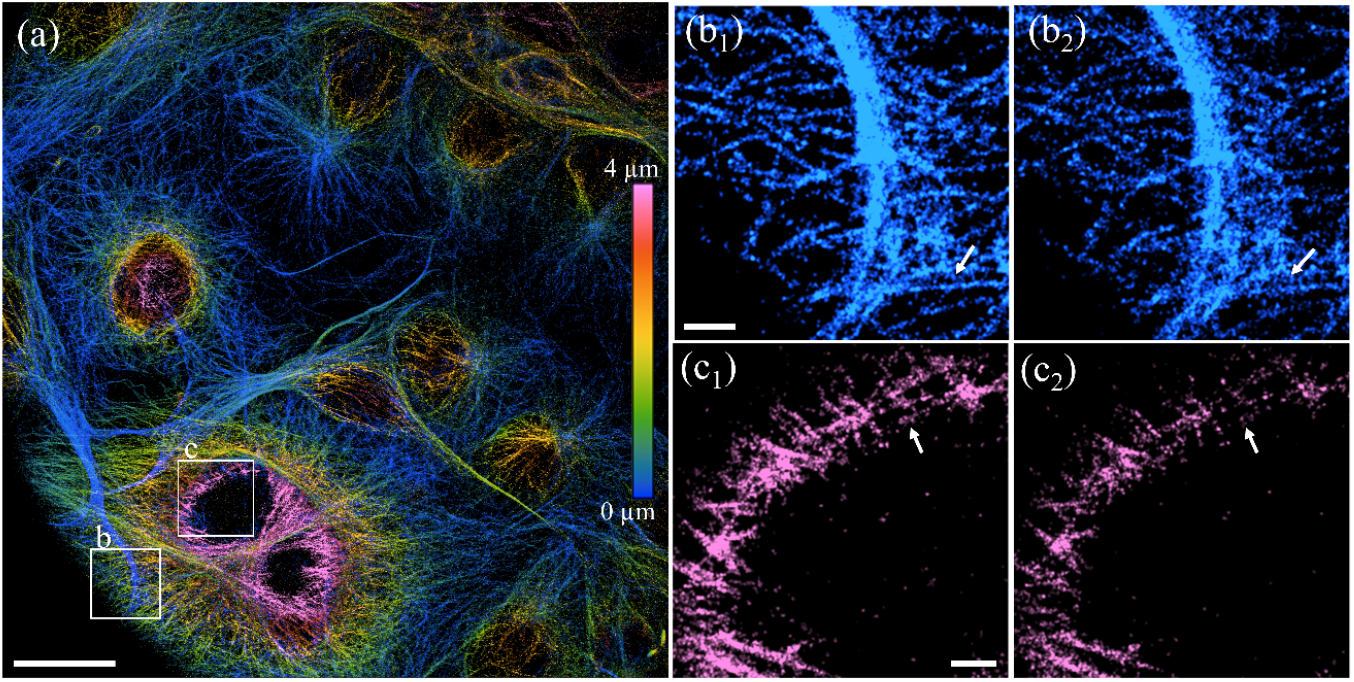
Large-FOV super-resolution image of microtubules. (a), microtubule image in a 130-by-131 μm FOV through FOV-dependent DeepSTORM3D. Scale bar: 20 μm. The reconstructed structure in square b at the depth of ∼100 nm is shown in (b_1_) (FOV-dependent-DeepSTORM3D) and (b_2_) (DeepSTORM3D). The structure in square c at the depth of 3900 nm is similarly shown in (c_1_) and (c_2_). Scale bar: 2 μm.

## 3. Discussion

In this work we demonstrate a field-dependent dense localization method for 3D SMLM. The method consists of two main components. First, we introduce PPG3D, a spatially variant PSF generator that employs PCA-based interpolation. This approach involves transforming PSFs into a lower-dimensional space, conducting interpolation for a target spatial position, and projecting back to the image space to obtain the desired PSF. Comparative analysis against other PSF generators in LM showcases significant enhancements in accuracy and operation speed. Then, we integrate PPG3D with DeepSTORM3D to address field dependence in large-FOV 3D LM, resulting in FOV-dependent-DeepSTORM3D. This new localization estimator demonstrates an improvement in prediction accuracy by over 30%. In STORM experiments, FOV-dependent DeepSTORM3D enables 3D super-resolution imaging of mitochondria and microtubules within a large FOV, exhibiting noticeably higher fidelity compared to standard DeepSTORM3D.

PPG3D offers a fundamental advantage in its simplicity, stemming from its interpolation nature, compared to model-based PSF generators. Reliable implementation of model-based generators often requires users to accurately retrieve physical parameters and comprehend physical models. In contrast, the PPG3D codes we provide only require calibrated PSFs and positions as input to accurately conduct PSF generation, making it potentially more user-friendly. Furthermore, the dimensionality reduction achieved through PCA confers a significant speed advantage to PPG3D, as it involves interpolating a smaller number of parameters. When compared to the shift variant ZPPR, PPG3D not only demonstrates superior accuracy and speed but also requires significantly fewer calibration positions.

While PSF generators based on decomposition and interpolation can be found in other fields, such as astronomy and computational volumetric imaging, PPG3D distinguishes itself from them in two key aspects. Firstly, PPG3D stands out as a genuine 3D continuous-domain PSF generator. In contrast, shift-variant PSF generation in astronomy^28^ is primarily conducted in 2D. Although volumetric imaging^29,30^ requires a 3D PSF, the voxel representation of objects dictates that the PSF remains unchanged within each discrete 3D voxel. Secondly, PPG3D employs a concise and efficient scheme of local interpolations, allowing for high-speed operation without any concerns regarding memory limitations. In comparison, other studies^29,30^ often resort to decomposing all the calibrated PSFs, which can pose challenges regarding memory efficiency.

However, the absence of a physical model limits PPG3D when the influence of refractive index mismatch is prominent. This limitation can be addressed, by combining a shift-invariant model-based PSF generator with PPG3D. For FOV-dependent-DeepSTORM3D, it enables field dependence learning, however cropping the whole FOV into many fixed sub-areas in advance is still naïve and lacks flexibility. A possible future direction can be to improve the flexibility of our method e.g. by randomly sampling the sub-images in the training stage and ultimately alleviating the need for positional encoding.

Finally, the combination of a field-dependent PSF generator with a deep learning-based localizer, presented here, can be used in a variety of scenarios, including where the PSF is controllably different throughout the FOV^32^.

## 4. Methods and Materials

### 4.1 Formulation of spatially variant PSF

The PSF is the impulse response of a system, which, in the imaging context, is the 3D electromagnetic field distribution at the image domain, in response to a point source at the object domain. For shift-invariant imaging systems, the PSF can be described, up to scaling and inversion, by:

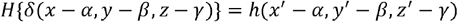

where *H* is the 3D transfer function, *δ* is the delta function, (x, y, z) are spatial coordinates of the object side and (x’, y’, z’) are for the image side, (*α,β,γ*) is the spatial position of the point source, and *h* is the 3D shape of the PSF. Note that we ignore the magnification effect for simplicity. If we abandon the shift-invariant assumption and consider the more general case, the PSF formation process can be given by:

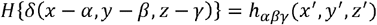

where now PSF *h*_*αβγ*_ is dependent on the location of the point source and the PSF shape changes as we move the point source in 3D object space. In practice, cameras with 2D sensors are typically used in imaging systems to sample the 3D space in a 2D plane. Switching to a 2D case, the impulse response becomes:

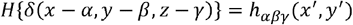

The PSF most obviously changes with the axial coordinate y, related to the concepts of “focus”, “defocus”, and “depth of field (DOF)”. In comparison, *αpβ* dependence is much milder and is typically ignored, for the sake of simple calculations. Generally, the former is called depth dependence and the latter is called field dependence, and the systems exhibiting field dependence are shift-variant. Our goal is to build a PSF generator that considers both kinds of dependence, for LM.

The basic idea of PPG3D is to transform the calibrated PSFs to a space with reduced dimensionality through PCA, specifically truncated singular value decomposition, and then implement interpolation in this low-dimensional domain with respect to the spatial position (*α,β,γ*). It is worth mentioning that, under this concept, different PCA interpolation schemes can be designed, dependent on the specific goal.

### 4.2 Imaging system calibration

To conduct the calibration experiment in a large FOV, two factors should be considered: point source distribution throughout the FOV, and homogeneous illumination. In this work, we use both fluorescent bead samples and a nano-hole array (Table S2 in SI) and rely on axial sample scanning by a microscope stage. For illumination, we built an illumination system composed of a 2-W high-power laser, a multi-mode fiber, and a vibration motor. Notably, SMLM experiment often benefits from strong illumination to enable efficient biomolecule blinking.

In the imaging system used for the STORM experiment, PSF engineering is performed using a Tetrapod diffractive optical element ^7^. We use a fluorescent bead sample immersed in water for calibration. After many times of lateral scanning, 334 field positions are covered within a circular FOV with a diameter of ∼180 μm. At each field position, the PSF is measured in an axial range of (−3, 3) μm at an interval of 0.15 μm. Thus, in total, this calibration consists of 334-by-41 spatial positions.

Before feeding this PSF dataset to PPG3D, we need to address the refractive index (RI) mismatch issue, i.e. the RI difference between the sample medium and the objective immersion oil. The image of a fluorescent source (e.g. sub-diffraction bead) on a glass coverslip is different from that of a fluorescent source inside a sample (Fig. S6 of SI); this is why the PSF derived by straight-forward interpolation of a z-stack of a bead on a glass coverslip cannot be used directly to obtain the PSF of a defocused emitter inside practically any biological medium at high accuracy.

Theoretically, the existence of the RI mismatch introduces two axial parameters with different contributions to the phase at BFP: the distance of an emitter from the coverslip, and the position of the nominal focal plane (NFP), i.e. the focal plane without RI mismatch. In calibration, we can only change the NFP positions, while the real experiment has fixed NFP and unknown various emitter positions. To transform the calibration measurements to be applicable to this RI case, we use model-based VIPR^18^. Specifically, we rely on the imaging model from VIPR to search for a proper NFP position such that the lowest emitter in experiment can be covered by the model. Then we fix the NFP and generate PSFs for the emitter at a series of distances relative to the coverslip. By doing so, we obtain a new PSF dataset with 334 field positions in a 180 μm-diameter FOV and 28 axial positions over a 4 μm range. PPG3D based on this dataset can now be used to generate shift-variant PSFs of emitters.

### 4.3 Sample preparation for STORM experiment

22×22 mm, 170 μm cover glasses (Deckgläser, No.1.5H) were cleaned in an ultrasonic bath with 5% Contrad 70 (Decon) at 60oC for 30 min, then washed twice with double distilled water (incubated shaking for 10 min each time), incubated shaking in ethanol absolute for 30 min, sterilized with filtered 70% ethanol for 30 min and dried in a biological cabinet. COS7 cells at a concentration of 60,000 cells/ml in Dulbecco’s Modified Eagle Medium (DMEM) with 1g/l D-glucose (Sartorius, 01-050-1A), supplemented with fetal bovine serum (Biological Industries, 04-007-1A), penicillin-streptomycin (Biological Industries, 03-031-1B) and glutamine (Biological Industries, 03-020-1B), were grown for 24 hr in a 6-well plate (Thermo Fisher, Nunclon Delta Surface) containing 6 ml of the cell suspension and the cleaned cover glasses, at 37oC, and 5% CO2. The cells were fixed with 4% paraformaldehyde and 0.2% glutaraldehyde in PBS, pH 6.2, for 60 min, washed and incubated in 0.3M glycine/PBS solution for 10 minutes. The cover glasses were transferred into a clean 6-well plate and incubated in a blocking solution for 2 hr (10% goat serum, 3% BSA, 2.2% glycine, and 0.1% Triton-X in PBS, filtered with 0.45 um Millex PVDF filter unit). The cells were then immune-stained with either 1:500 diluted anti-TOMM20-AF647 antibody (Abcam, ab209606) or 1:500 diluted anti-alpha-tubulin-AF647 (ab190573) and 1:500 diluted anti-beta-tubulin-AF647 (Abcam, ab235759) in the blocking buffer for 1.5 hr and washed five times with PBS. For super-resolution imaging, a PDMS chamber (22×22×3 mm, with a 13×13 mm hole cut in the middle) was attached to the cover glass containing the fixed and stained COS7 cells to create a pool for the blinking buffer. Blinking buffer (50 mM Cysteamine hydrochloride (Sigma, M6500), 20% sodium lactate solution (Sigma, L1375), and 3% OxyFluor (Sigma, SAE0059) in PBS, pH 8-8.5) was added and a cover glass was placed on top while ensuring minimal air bubbles.

### 4.4 STORM Imaging

We used the Nikon eclipse Ti2 inverted microscope equipped with N-STORM unit (Nikon), silicone-oil objective (Nikon, RS HP Plan Apo 100x/1.35 Sil WD), and a multi-bandpass dichroic (Semrock, Di03-R405-488-532-635-t3). The microscope was extended with a 4f system (f=200 mm) containing a tetrapod phase mask in the Fourier plane and a sCMOS camera (Teledyne Photometrics, Kinetix) for image acquisition. The sample was illuminated by a 640 nm, laser at estimated power of ∼3.13 kW/cm^2 at the sample.

## Supporting information

Reconstruction of mitochondria

Reconstruction of microtubules

## 5. Acknowledement

We would like to express our sincere gratitude to Ofri Goldenberg for fruitful discussions and Shuang Fu for his assistance with the shift-vairant ZPPR codes. This research was supported in part by the ISRAEL SCIENCE FOUNDATION (grant No. 450/18); and by funding from the European Union’s Horizon 2020 research and innovation program under grant agreement No. 802567 -ERC-Five-Dimensional Localization Microscopy for Sub-Cellular Dynamics. YS is supported by the Zuckerman Foundation.

## 6. Codes

PPG3D: https://github.com/dafeixiao/PPG3D

FOV-dependent DeepSTORM3D: https://github.com/dafeixiao/FOV-dependent-DeepSTORM3D

