## Supplementary material for "Large-FOV 3D localization microscopy by spatially variant point spread function generation": SI

**Supplementary information**

1. **Optical imaging systems**

The imaging systems involved in this work can be illustrated as Fig. S1 and their detailed compositions and parameters are presented in Table S1. The samples we used for calibration and super-resolution imaging tasks are described in Table S2. For PPG3D’s demonstration and comparisons (Fig. 2 of main text) with other point spread function (PSF) generators in LM, the image system # 1 with an oil objective (1.35 NA, 100 magnification) and bead samples are used. For the field dependence quantification (Fig. 4-5), we used the image system # 1 with an air objective (0.75 NA, 40 magnification) and the nanohole array. The SOTRM experiments were implemented in imaging system # 2 with the oil objective (1.35 NA, 100 magnification) and bead samples were used for calibration.

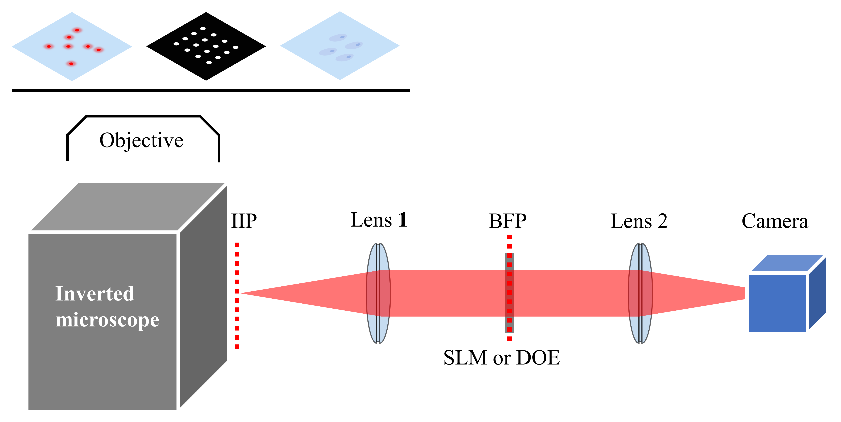

Fig. S1. Optical imaging system illustration. The imaging system is composed of an inverted Nikon microscope and an 4f optical extension with a phase modulator (spatial light modulator, SLM for short, or diffractive optical element, DOE for short) at the back focal plane (BFP) for Tetrapod PSF engineering. IIP denotes intermediate image plane. Bead samples, a nanohole array and cells with fluorescently labeled mitochondria or microtubules were used.

Table S1. Compositions and parameters of imaging systems

| Optical  imaging system | Microscope | 4-f setup | | |
| --- | --- | --- | --- | --- |
|  |  | Focal length of both lenses | Phase modulator | Camera |
| # 1 | Nikon ECLEPSE Ti2 inverted microscope | 100 mm | SLM, Meadowlark, pixel size: 9.2 µm, resolution: 1152x1920 | Teledyne Photometrics Prime 95B, pixel size: 11 µm, resolution: 1200x1200 |
| # 2 |  | 200 mm | A Tetrapod DOE, fabricated by lithography | Teledyne Photometrics Kinetix, pixel size: 6.5 µm (2 times of binning are used), resolution: 3200x3200 |

Table S2. Details of samples

| Sample | Details |
| --- | --- |
| Beads (fluorescent microspheres) | - excitation/emission wavelength: 625/645 nm, diameter: 200 nm. - mixed with polyvinyl alcohol (PVA) solution, coated on a coverslip on a spin coater. - The medium of the sample is air (refractive index of 1.0) or water (refractive index of 1.33) by dropping distilled water above. |
| Nano-hole array | - consist of 50-by-50 holes with a diameter of 200 nm and an interval of 20 µm from each other. - fabricated on a coverslip with a chrome coating of 100 nm thickness, drilled by ion beam etching (IBE). |
| Cells | - label mitochondria or microtubules. - Labeling details in Methods of the main text. |

2. **Processing image stacks from calibration experiments**

The shift-invariant mode or axial mode of PPG3D simply receives a PSF stack with the corresponding axial positions. For shift-variant mode, we need to input a package of 4-dimensional PSF dataset, a list of lateral positions and another list of axial positions.

The dimensions of the PSF dataset are the number of lateral positions, axial positions, and pixels in height and width. To crop PSFs from the calibrated images, PSF center detection is required and we resort to cross-correlation and template matching to achieve that. With an example of processing one image stack, we follow four steps.

1) choose one image (superposition of many PSFs at one axial plane) from the stack and manually crop one PSF as the template.
2) calculate the cross correlation of the cropped PSF template with the chosen image.
3) search for peaks of the correlation result. Peak detection without the prior knowledge of the peak number is not a seemingly easy task. Here we first resort to a coarse but fast algorithm [1] which can reach the approximate region around the real peaks, and then apply Gaussian fitting near the approximately searched peaks to find a fine detection with a sub-pixel accuracy.
4) crop the image stack according to the searched PSF centers and form the 4D PSF dataset.

After processing all the image stacks, we filter out field positions very close to each other. Finally, we combine the valid field positions and axial positions and obtain the input of PPG3D.

3. **PPG3D implementation details**

As mentioned in the main text, for one PSF interpolation in 3D, we use three calibrated field positions, two axial positions, and total six spatial calibration positions. The searching criterion regarding those positions mentioned in the main text consists of the following procedures:

1) calculate lateral distances of the target position from all the calibrated positions.
2) choose a certain number of closest positions from which we combine any three of them to form a triangle.
3) calculate the centroid of each triangle.
4) choose the centroid closest to the target position and the corresponding three field positions are the final selection (see Fig. S2 (a)).
5) the two axial planes are chosen simply in terms of axial distance.

One example about the field position selection and interpolation is shown in Fig. S2.

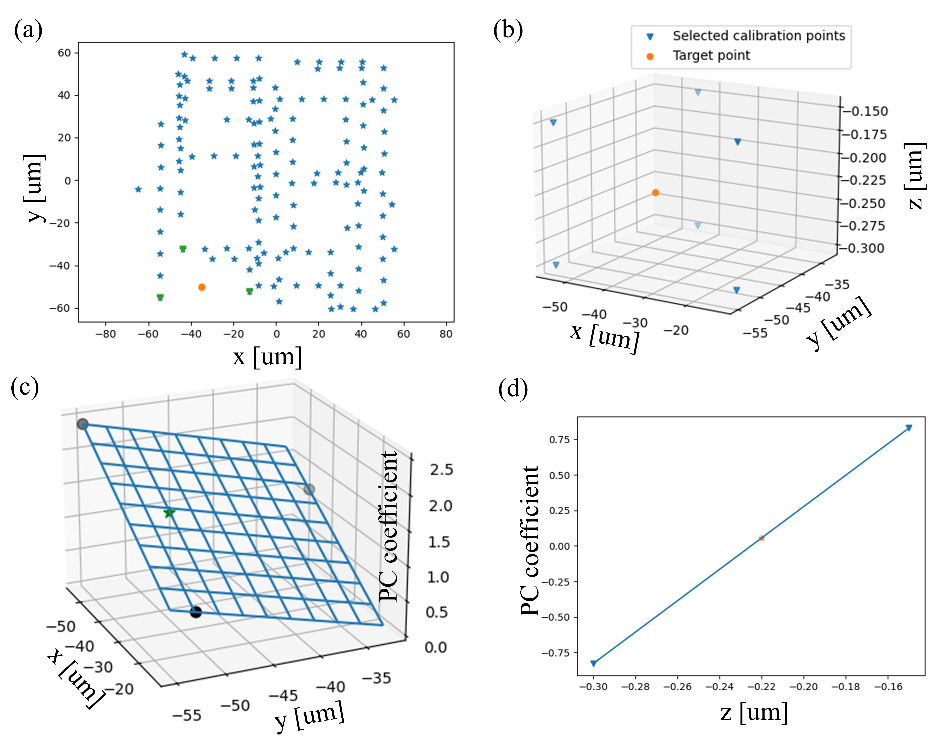

Fig. S2. PPG3D interpolation details. (a), three selected field positions (green) around the target position (orange). (b), six searched positions (blue) around the target position (orange). (c), an example of one 2D lateral interpolation regarding principle component coefficients where the green star dot from interpolation. (d), an example of one 1D axial interpolation where the orange star point is from interpolation.

4. **VIPR and ZPPR implementations**

VIPR [2] aims at retrieving a pixel-wise phase mask at BFP to characterize the deviations of a practical imaging system from the ideal imaging model (scalar or vectorial). ZPPR, short for Zernike-polynomial pupil phase retrieval, differs from the VIPR concept in the pupil formulation. For the imaging system # 1 in table S1, the retrieved phases from both methods are shown in Fig. S3 (a) and (b). When it comes to shift-variant ZPPR, the regular ZPPR is implemented to yield one vector of Zernike coefficients at each calibrated field position from which vectors at any other non-calibrated positions are interpolated. Based on data in the Fig. 2 of the main text, one example of such interpolation is shown in Fig. S3 (c). Specifically, 146-by-21 Zernike coefficients are retrieved at 146 field positions and another 5 “unseen” positions (marked as red cross dot in Fig. S3 (c)) are for test.

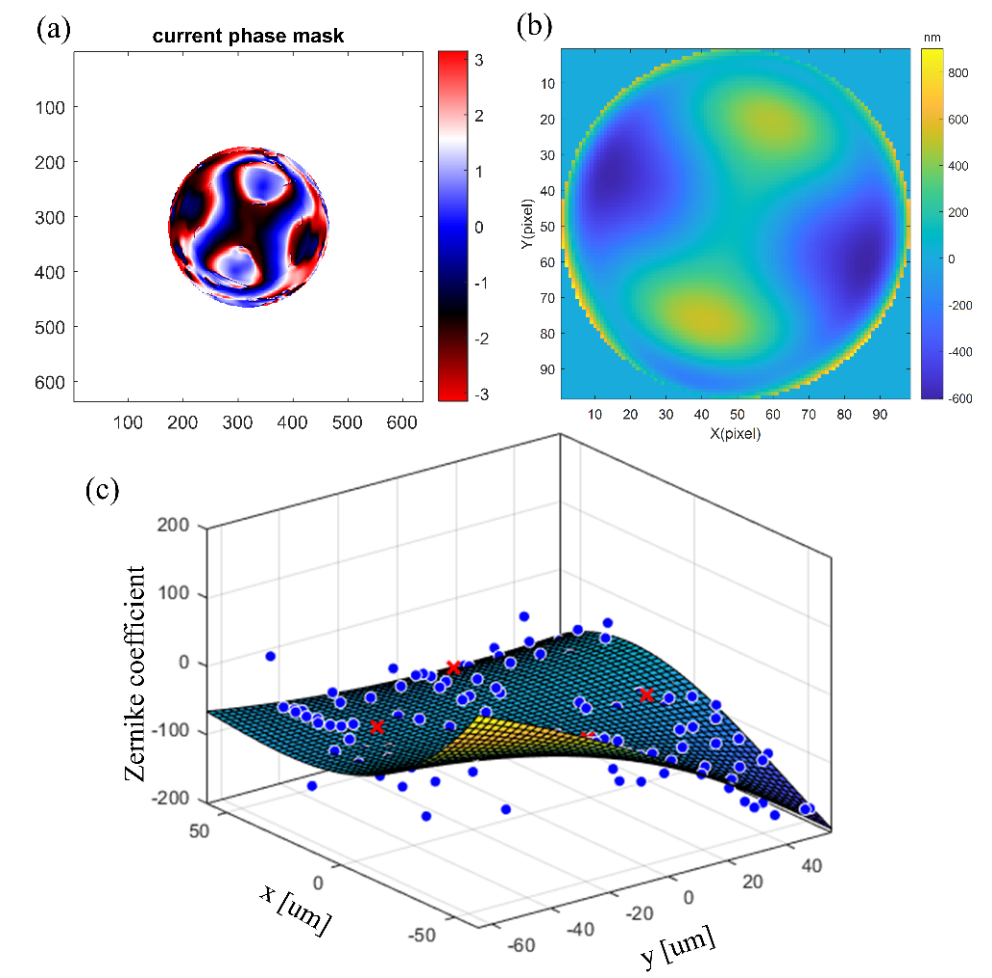

Fig. S3. (a), The retrieved phase from VIPR at P0 position in the Fig. 2 (a) of the main text. (b), retrieved phase from ZPPR with 21 Zernike basis functions at the same P0 position. (c), an example of 2D interpolation of Zernike coefficients in shift-variant ZPPR implementation. The blue dots and red cross dots are the “seen” calibration positions and “unseen” test positions respectively.

5. **Field dependence quantification**

The utilization of PCA on calibrated PSFs can also serve to quantitatively characterize the field dependence of an imaging system. Here, we use a nano-hole array for calibration in an imaging system (optical setup #1 of Table S1) with a 40x objective. We captured PSFs at 234 field positions (Fig. S4(a)) within an axial range of (-3 µm, 3 µm), with intervals of 0.2 µm. To quantify the difference of these PSFs relative to a central reference PSF at a specific axial plane, we employ PCA-based Euclidean distance, which measures the vector distance in PCA space.

Initially, we computed the Euclidian distance and obtained non-circular field dependence, shown as a contour plot (Fig. S4 (b)). This plot reveals that the PSF changes more rapidly towards the right side of the (FOV) (as indicated by the higher density of contour lines on the right). Additionally, the measured PSF, at the right red spot below the contour plot, appears to be significantly defocused, which can be attributed to a slight tilt of the object slide or the microscope stage—a common phenomenon encountered in large FOVs.

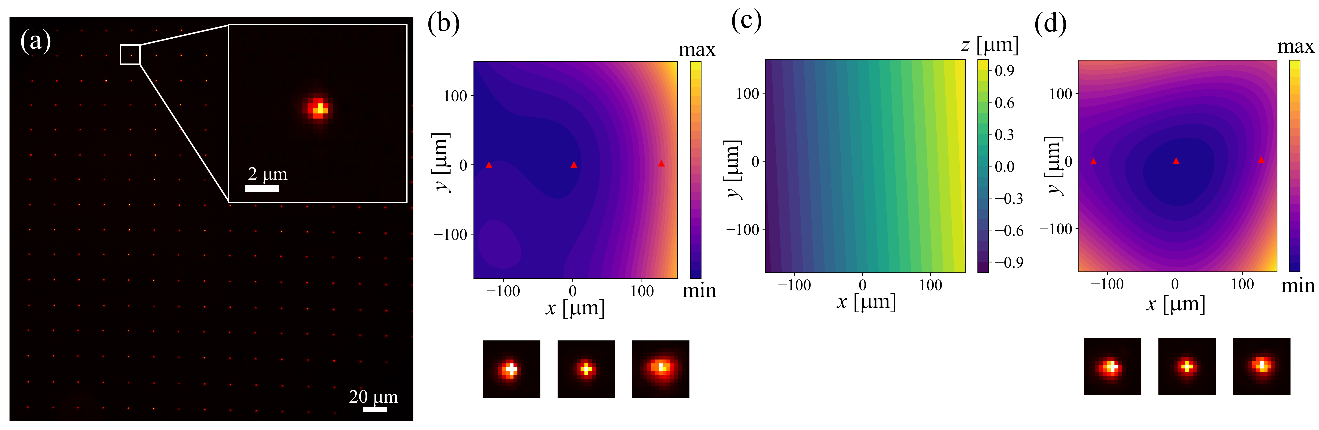

Fig. S4. Field dependence characterization with tilt detection. (a) an image of a nano-hole array. (b) characterization through PCA-based distance without tilt compensation. Three PSFs at the red spots are shown below. (c) Contour of the detected tilt plane with direction cosine of -0.0063 along x and -0.0003 along y. (d) field dependence characterization with tilt compensation where three PSFs at the red spots (same as (b)) are shown below.

Significantly, the tilt effect often exhibits a more pronounced field-dependent impact compared to other sources of field dependence in an imaging system, such as aberrations introduced by the objective lens or the PSF engineering module. To mitigate the influence of tilt, we employ a four-step method for tilt detection:

1) Choose a reference field position.

2) Utilize PPG3D to axially interpolate PSFs at other field positions.

3) At each field position, identify the axial position where the PSF most closely resembles the reference PSF.

4) Fit all searched axial positions to a tilted plane.

The detected tilt (Fig. S4 (c)) demonstrates a maximum axial position difference of ~2 µm. After compensating for the tilt effect by subtracting the calculated tilt value at each calibrated position, we reevaluated the field dependence (Fig. S4 (d)), which now appears circularly distributed. The corrected PSFs at the same three field positions, as shown in Fig. S4 (d) after tilt compensation, demonstrate reduced bias in field dependence compared to those before compensation (Fig. S4(b)).

The advantage of quantifying field dependence in PCA space is robustness. Because the bases of PCA is from all the calibrated PSF data, the vector distance in PCA space reflects global difference, making it less dependent to noise. However, it becomes unconvincing to compare vectors in different PCA spaces. To quantify field dependence in different imaging systems (with and without PSF engineering), we resort to correlation coefficient (CC).

We use PSF dataset in Fig. 2. of main text, as well as the corresponding dataset without PSF engineering (switching off the tetrapod phase mask at the SLM), to demonstrate the influence of PSF engineering on field dependence. As the implementation before, we first detect the tilt of the same imaging system (# 2 of Table S1) without (Fig. S5 (b)) and with (Fig. S5 (d)) PSF engineering. This yields consistent results: the direction cosine along x and y axis of the tilt norm in (b) is (0.0020, 0.0030), and that in (d) is (0.0020, 0.0033)).

With the tilt compensation, we obtain both field dependence quantification results (Fig. S5 (c) and (e)). The smaller CC value in (e) reveals that field dependence is exacerbated due to PSF engineering, which might be caused by practical factors, e.g., the placement and operation error of the SLM.

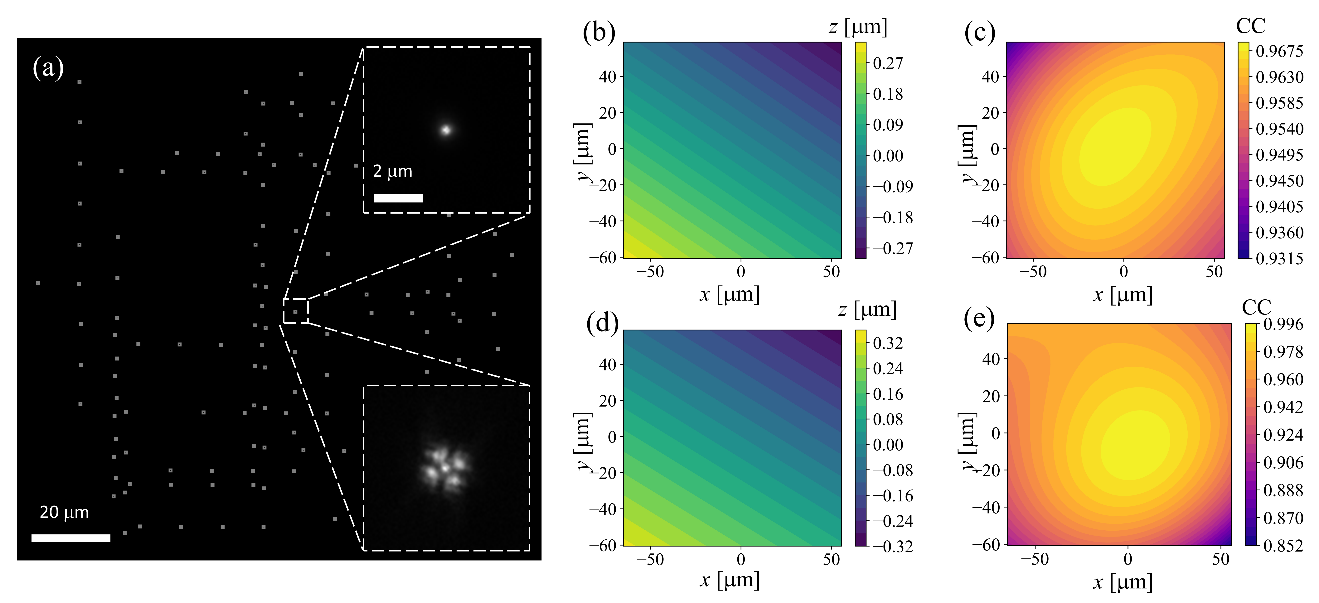

Fig. S5. Influence of PSF engineering on field dependence quantification of an imaging system. (a), FOV of the imaging system (the same as Fig. 2 (a) in main text) with calibrated field positions. The zoom-in view shows the PSF in two cases: without and with PSF engineering. The former yields tilt detection (b) and the field dependence quantification result (c) through CC metric and the latter yields (d) and (e).

6. **Refractive index mismatch**

When refractive index (RI) mismatch exists (Fig. S6 (a)), one needs to consider two axial positions to determine the defocus phase at pupil: the distance of the emitter above the coverslip ($\gamma_{e}$) and the distance of the nominal focal plane (NFP) position above the coverslip ($\gamma_{NFP}$). We assume that a positive position is above the coverslip. These two axial parameters have different contributions to the spherical defocus phase at the BFP. The electric field at BFP can be formulated as

$U_{BFP}=circ(\rho)exp(j(M+\gamma_{e}k_{2}\sqrt{1-{(\frac{n_{1}}{n_{2}}\rho)}^{2}}-\gamma_{NFP}k_{1}\sqrt{1-\rho^{2}}+2\pi(\alpha\frac{xA_{M}}{\lambda f_{4f}}+\beta\frac{yA_{M}}{\lambda f_{4f}})))$---(S1)

where

$\rho=\frac{\sqrt{x^{2}+y^{2}}}{r_{p}}\frac{\mathrm{NA}}{n_{1}}$ ---(S2)

$r_{p}=\frac{f_{4f}NA}{\sqrt{{A_{M}}^{2}-{NA}^{2}}}$---(S3)

and description of the involved parameters is shown in Table S3. The different contributions can be revealed from the different multiplication factor before $\gamma_{e}$ and $\gamma_{NFP}$. In the case without RI mismatch, namely, n_1_=n_2_, k_1_=k_2_, these two factors are the same and can be combined according to the associative law, yielding the fact that the spherical defocus phase merely depends on the relative position of the emitter with respect to the NFP.

Table S3. Parameter table

| Parameter | Description |
| --- | --- |
| ($x$, $y$) | Cartesian coordinates at BFP |
| $\rho$ | Normalized polar coordinate at BFP |
| ($\alpha, \beta, \gamma_{e}$) | Cartesian coordinates of an emitter, $\gamma_{e}$ is the relative distance above the coverslip |
| $\gamma_{NFP}$ | The distance of the nominal focal plane above the coverslip |
| M | Phase mask at BFP |
| $A_{M}$ | Magnification of the objective |
| $n_{1}, k_{1},n_{2}, k_{2}$ | Refractive index of the medium between coverslip and objective, corresponding wave number, refractive index of the medium inside the sample, corresponding wave number |
| $\lambda$ | Wave length of the emission light |
| $f_{4f}$ | Focal length of the first lens in the 4-f setup |
| $circ(\rho)$ | A circular window with 1 inside and 0 outside. The window radius is NA/n_1_ for oil objective and supercritical angle fluorescence can be included |

To visualize the RI mismatch influence on PSF, we apply the scalar imaging model (imaging system # 1 of Table S1 where n_1_ and n_2_ are 1.406 and 1.0 respectively) and compare PSFs at ($\gamma_{e}, \gamma_{NFP}$) and ($\gamma_{e}+0.5, \gamma_{NFP}+0.5$). Both PSFs should be the same if there is no such RI mismatch, while this resemblance is broken in mismatch case (Fig. S6 (c)). In order to transform PSFs at ($\alpha, \beta,\gamma_{NFP}$) from the calibration experiments to PSFs at practical ($\alpha, \beta,\gamma_{e}$), we rely on the physical imaging model embedded in VIPR [2].

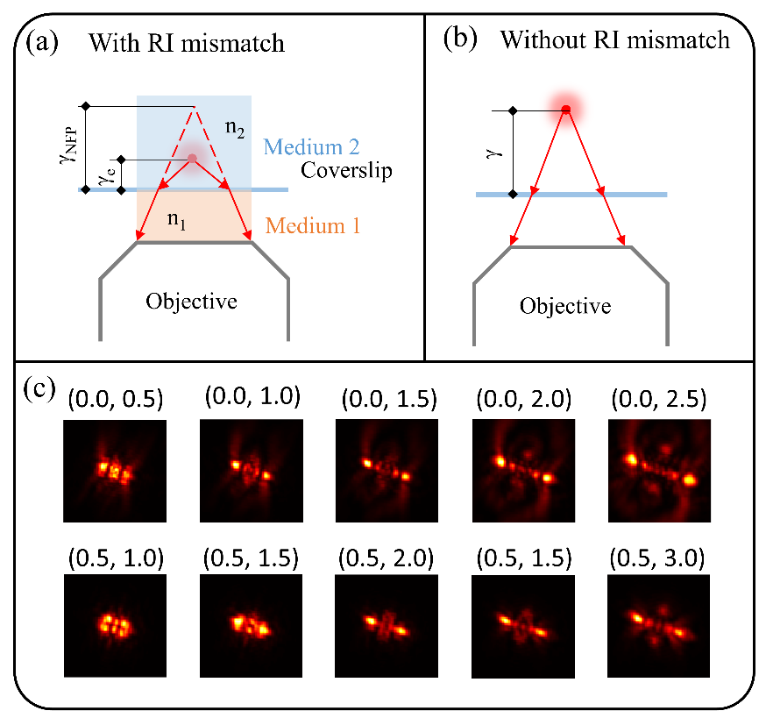

Fig. S6. (a) and (b), Illustration of RI mismatch in microscopy. (c), influence of RI mismatch on PSFs at a series of axial pairs ($\gamma_{e}, \gamma_{NFP}$).

7. **SMLM simulation and experiment details**

**7.1. FOV specification**

When it comes to a large FOV or high throughput, imaging tasks takes a lot of time. For the FOV-dependent DeepSTORM3D, as a variant of DeepSTORM3D with additional input layers of transverse coordinate maps, it takes several days in training (90000 training images and 10000 test images for both networks) and several seconds in inference to localize emitters of one frame in our case (1371-by-1371 pixels in one frame, 11-by-11 squares of sub-images and 161 pixels in width and height of each sub-image, GPU: NVIDIA TITAN RTX with 24 G memory).

To demonstrate our concept in simulation, we think it is reasonable to focus on a FOV smaller than the calibrated large FOV for the sake of time saving. Specifically, our simulation is implemented in the blue area 1 (Fig. S7) which consists of a 6-by-6 grids of sub-areas with overlapping. Each sub-area has a valid central region (the central shade in Fig. S7) due to the consideration of PSF truncation at the edge. The “CCCC” micro structure (see the ground truth in Fig. S8) and the sinusoid micro surface (see the reconstructed result in Fig. S9) are set in the transparent blue and yellow region in Fig. S7 respectively. In experiments, we switch to the whole FOV (orange region 2 in Fig. S7).

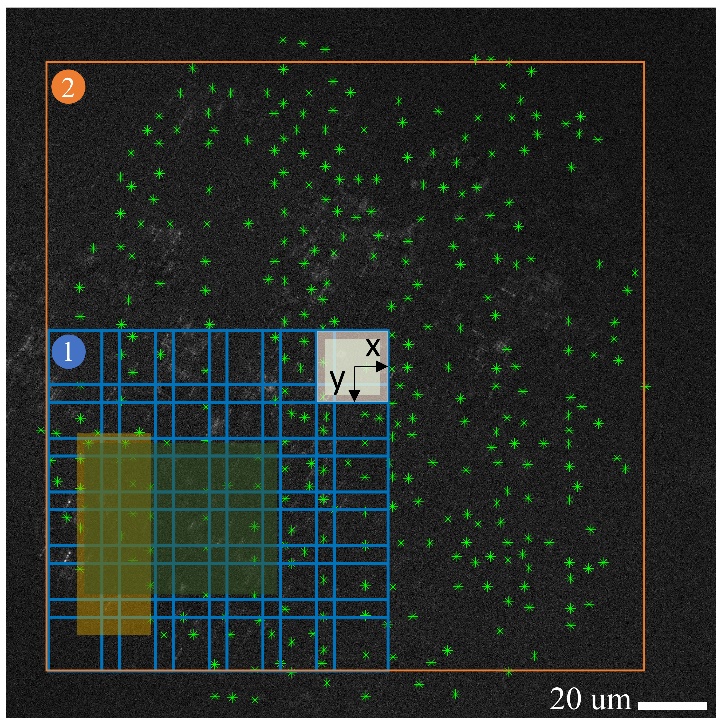

Fig. S7. Illustration of the calibration positions and FOV explanation in SMLM. The background is one example experimental image with green dots specifying calibration positions throughout the FOV. In simulation, the region 1 covered with 6-by-6 overlapping blue squares (sub image regions of 161 pixels) was used. The two shade squares at center presents one sub region and its valid central area (PSFs close to the edge are possibly truncated, thus are invalid). The blue shade rectangle with transparence is the area for “CCCC” simulation in the main text and the yellow one is for the sinusoidal surface simulation (Fig. S9). The orange rectangle region 2 shows the FOV in SMLM experiment with a width of 178 µm.

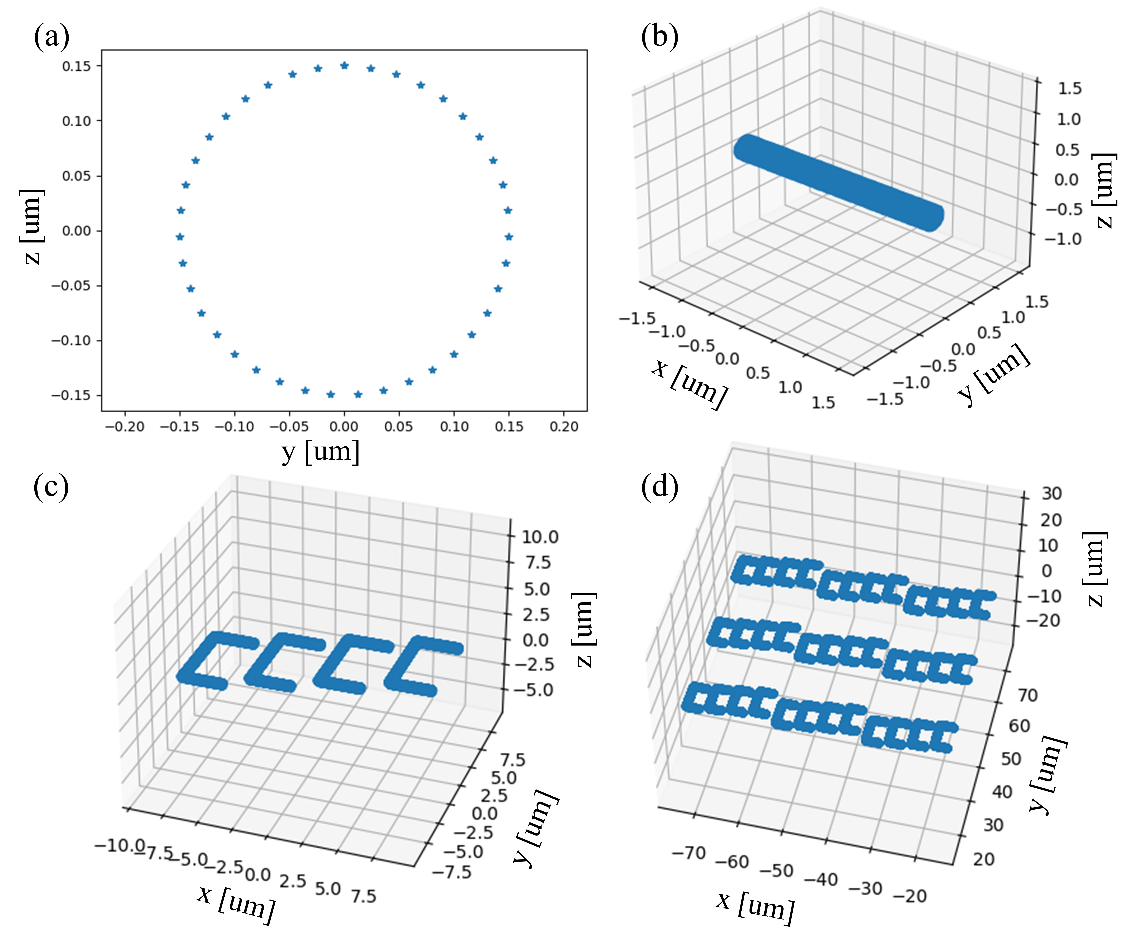

Fig. S8. Simulated dotted tubule “CCCC” structure. (a) is the cross section of the tubule unit (b). Each “C” includes four tubule units and the combination of the four letters is shown in (c). The axial locations of the four letters are 0.6 µm, 1.6 µm, 2.6 µm, and 3.6 respectively. The whole 3D micro structure (d) has 9 “CCCC” units.

7.2. **Additional simulation details and results**

The demonstration of the PPG3D’s contribution in SMLM is built upon a valid comparison between FOV-dependent DeepSTORM3D and regular DeepSTORM3D. Importantly, the localization numbers from both networks should be the same. As we mentioned in the main text, both networks yield different numbers of inference under the same post-processing threshold [3]. To address this difference, we resort to an inference confidence filter and density filter of ThunderSTORM (an ImageJ plugin) and finally obtain around 0.85 million valid predictions from both networks for the comparison of “CCCC” structure. This number is 0.35 million for sinusoid surface (Fig. S9).

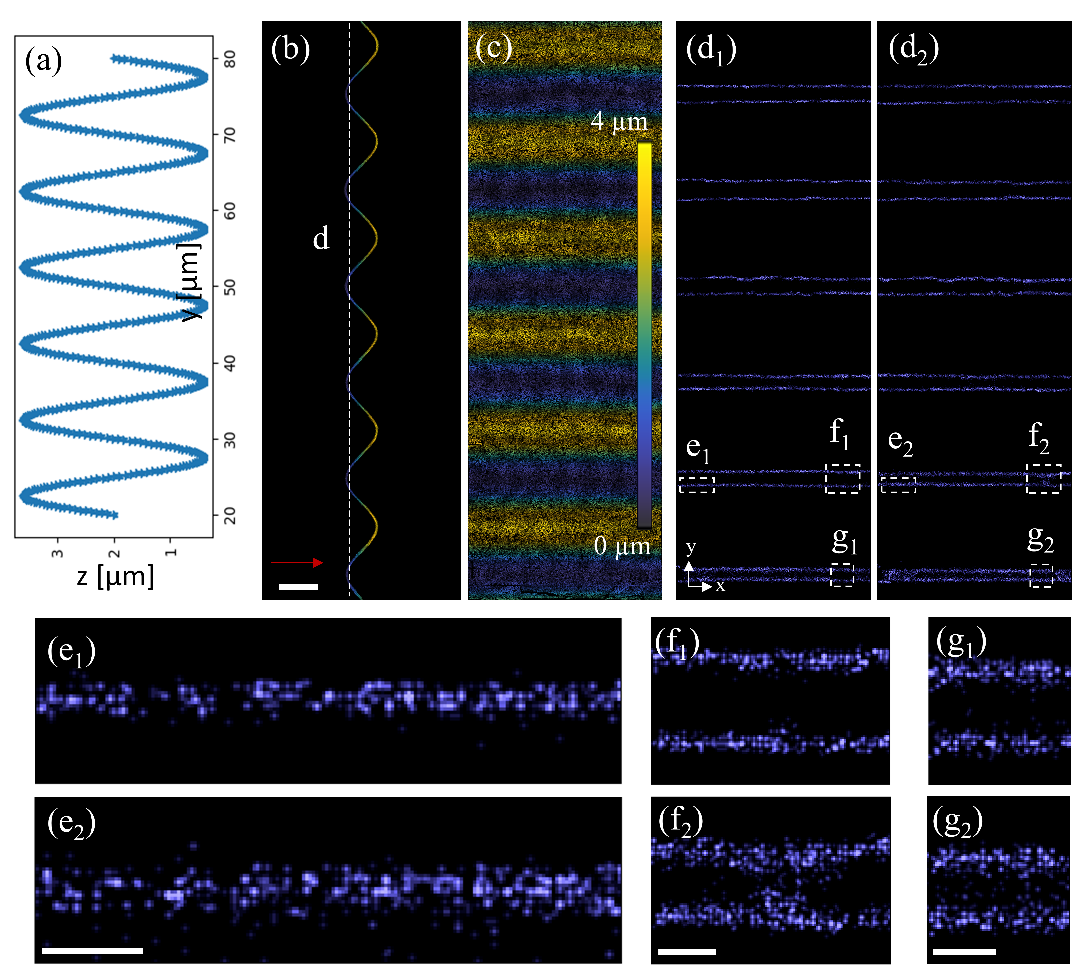

Fig. S9. Simulation of a sinusoidal surface in SMLM. (a), Ground truth of the surface’s cross section. (b), reconstructed surface, by FOV-dependent DeepSTORM3D, which is projected at the y-z plane (b) and x-y plane (c). The red arrow shows z axis. Scale bar: 4 µm. Cross section (d_1_ and d_2_) of the white line in (b) presents localization results by FOV-dependent DeepSTORM3D (subscript “1”) and DeepSTORM3D (subscript “2”). The zoom-in views of squares e, f, and g are shown in (e_1_), (e_2_), (f_1_), (f_2_), (g_1_), and (g_2_), which demonstrates improvements due to field dependence consideration.

7.3. **Additional experiment details and results**

In SMLM experiment, the number of valid localizations for the mitochondria (Fig. 6 in main text) is around 12.7 million and that number is 11.5 million for the microtubule reconstruction (Fig. 7 of main text) and 3.2 million for an additional microtubule imaging in Fig. S10.

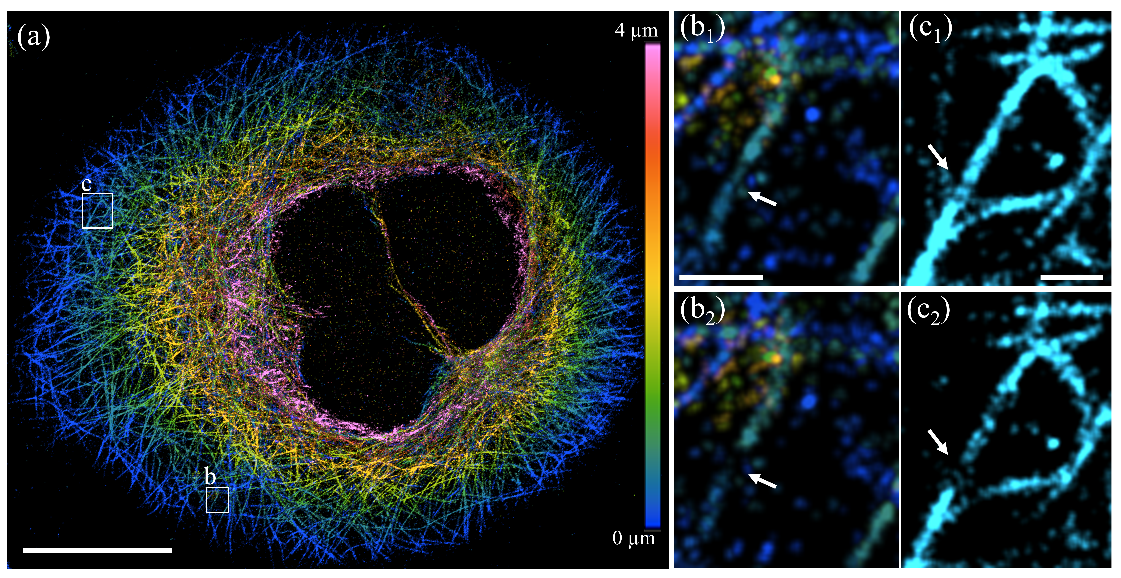

Fig. S10. An additional super-resolved imaging of microtubules. (a), microtubule image in a 130-by-131 µm FOV through FOV-dependent-DeepSTORM3D. Scale bar: 20 µm. The reconstructed structure in square b at the depth of 97.5 nm is shown in (b_1_) (FOV-dependent-DeepSTORM3D) and (b_2_) (DeepSTORM3D). The structure in square c at the depth of 3900 nm is similarly shown in (c_1_) and (c_2_). Scale bar: 1 µm.
